# Fluoxetine-induced dematuration of hippocampal neurons and adult cortical neurogenesis in the common marmoset

**DOI:** 10.1101/661025

**Authors:** Koji Ohira, Hideo Hagihara, Miki Miwa, Katsuki Nakamura, Tsuyoshi Miyakawa

**Affiliations:** Division of Systems Medical Science, Institute for Comprehensive Medical Science, Fujita Health University, Toyoake, Aichi 470-1192, Japan; Laboratory of Nutritional Brain Science, Department of Food Science and Nutrition, Mukogawa Women’s University, Nishinomiya, Hyogo 663-8558, Japan; Cognitive Neuroscience Section, Primate Research Institute, Kyoto University, Inuyama, Aichi 484-8506, Japan

## Abstract

The selective serotonin reuptake inhibitor fluoxetine (FLX) is widely used to treat depression and anxiety disorders. Chronic FLX treatment reportedly induces cellular responses in the brain, including increased adult hippocampal and cortical neurogenesis and reversal of neuron maturation in the hippocampus, amygdala, and cortex. However, because most previous studies have used rodent models, it remains unclear whether these FLX-induced changes occur in the primate brain. To evaluate the effects of FLX in the primate brain, we used immunohistological methods to assess neurogenesis and the expression of neuronal maturity markers following chronic FLX treatment (3 mg/kg/day for 4 weeks) in adult marmosets (n = 3 per group). We found increased expression of doublecortin and calretinin, markers of immature neurons, in the hippocampal dentate gyrus of FLX-treated marmosets. Further, FLX treatment reduced parvalbumin expression and the number of neurons with perineuronal nets, which indicate mature fast-spiking interneurons, in the hippocampus, but not in the amygdala or cerebral cortex. We also found that FLX treatment increased the generation of cortical interneurons; however, significant up-regulation of adult hippocampal neurogenesis was not observed in FLX-treated marmosets. These results suggest that dematuration of hippocampal neurons and increased cortical neurogenesis may play roles in FLX-induced effects and/or side effects. Our results are consistent with those of previous studies showing hippocampal dematuration and increased cortical neurogenesis in FLX-treated rodents. In contrast, FLX did not affect hippocampal neurogenesis or dematuration of interneurons in the amygdala and cerebral cortex.

## Introduction

The antidepressant fluoxetine (FLX) is one of the most widely used drugs for the treatment of depression and anxiety disorders. FLX is a selective serotonin reuptake inhibitor (SSRI) that prevents the reuptake of serotonin into presynaptic neurons [1], thereby maintaining increased serotonin levels in the synaptic region and promoting repeated stimulation of postsynaptic serotonin receptors [2]. Although SSRIs induce immediate increases in extracellular serotonin levels in the central nervous system (CNS), several weeks of treatment are typically required to elicit therapeutic effects [2]. Several adverse psychiatric effects of SSRIs also emerge after 2–3 weeks of chronic treatment or after treatment cessation [3,4]. Several changes have been reported during this time period in the CNS of animals treated with SSRIs. One of the most notable effects is an increase in adult hippocampal neurogenesis. Chronic FLX treatment (2−4 weeks) has been reported to increase neurogenesis in the adult dentate gyrus (DG) [5–7], a response that is critical to the behavioral effects of FLX [7]. Neural progenitor cells have been found in the adult cerebral cortex [8], and FLX treatment increases cortical adult neurogenesis from these progenitors [9].

Chronic FLX treatment also induces the reversal of neuronal maturation in the hippocampus, amygdala, and cerebral cortex. In the hippocampal DG of adult mice chronically treated with FLX, nearly all granule cells revert to a state featuring multiple molecular and electrophysiological characteristics of immature granule cells, a phenomenon termed “dematuration” [10]. FLX-induced dematuration is also observed in fast-spiking interneurons in the hippocampal CA1 and CA3 regions, amygdala, and medial frontal cortex, as indicated by significant decreases in parvalbumin (PV) expression and perineuronal nets (PNNs), markers of mature fast-spiking interneurons [11,12]. We recently showed that genome-wide gene expression patterns in the DG and frontal cortex of adult mice chronically treated with FLX were similar to those of typically developing infant mice [13], supporting the idea that FLX induces dematuration in the brain regions in terms of transcriptomic level.

These molecular and cellular changes induced by chronic FLX treatment may be involved in mechanisms underlying both the therapeutic and adverse effects of SSRIs. However, evidence of these changes has been mainly derived from rodent models, and it remains unclear whether FLX treatment induces similar effects in primates. Therefore, the aim of this study was to use immunohistological methods to determine whether and to what extent chronic FLX treatment alters neuron maturity and neurogenesis in the marmoset brain.

## Materials and Methods

### Animals

Six experimentally naïve male common marmosets (*Callithrix jacchus*) of ages ranging from 2–8 years were used in this study. Animals were kept at the Primate Research Institute of Kyoto University. Every effort was made to minimize the number of animals used.

### FLX treatment

The animals were divided into 2 groups. One group was treated with FLX pellets (Innovative Research of America, Sarasota, FL) for 4 weeks at a dose of 3 mg/kg/day, whereas the other group was treated with control pellets (Innovative Research of America) for the same period. This dosage was chosen based on results of a previous study showing behavioral alterations in marmosets receiving the same dose [14]. Animals were anesthetized with an intramuscular injection of ketamine hydrochloride (50-60 mg/kg) and medetomidine hydrochloride (0.1-0.15 mg/kg), followed by atipamezole hydrochloride injection of 0.5-0.75 mg/kg. FLX or control pellets were implanted subcutaneously on their backs [15].

### Bromodeoxyuridine labeling

Bromodeoxyuridine (BrdU) injections were performed as previously described [16]. Briefly, a stock solution of 20 mg/mL BrdU (Sigma-Aldrich, St. Louis, MO, USA) in distilled water with 0.007 N NaOH was prepared and stored at −20°C until use. Animals were intraperitoneally injected with BrdU diluted in phosphate-buffered saline (PBS; 100 mg/kg) every 24 h for 3 days starting 14 days after FLX treatment onset.

### Immunohistological analysis

Animals were deeply anesthetized with pentobarbital sodium (20 mg/kg) and medetomidine (0.02 mg/kg) and transcardially perfused with 4% paraformaldehyde in 0.1 M phosphate buffer, pH 7.4. The brains were removed, immersed in 4% paraformaldehyde overnight at 4°C, and transferred to 30% sucrose in PBS for at least 7 days for cryoprotection. Brain samples were mounted in Tissue-Tek (Miles, Elkhart, IN), frozen, and cut into 50 μm-thick coronal sections using a microtome (CM1850; Leica Microsystems, Wetzlar, Germany). Sections were stored in PBS containing sodium azide (0.05%, w/v) at 4°C until use.

BrdU staining was performed as previously described [16]. Briefly, sections were incubated at 4°C for 10 min in 0.1 N HCl and then at 37°C for 30 min in 2 N HCl. Sections were washed twice for 5 min in PBS and then blocked in 0.2 M glycine in PBS at room temperature for at least 2 h. The following procedures were the same as methods with other primary antibodies.

After washing in PBS for 1 h, sections were preincubated with PBS-DB (4% normal donkey serum [Vector Laboratories, Burlingame, CA] and 1% bovine serum albumin in PBS) for 2 h at room temperature. The sections were then incubated at 4°C for 48 h or at room temperature overnight with primary antibodies. After washing in PBS for 1 h, the sections were incubated at room temperature for 1 h with secondary antibodies. Sections were then washed in PBS containing Hoechst 33258 (Sigma-Aldrich) for 1 h to counterstain nuclei, mounted on glass slides coated with 3-aminopropyltriethoxysilane, and embedded with PermaFluor (Thermo Fisher Scientific, Waltham, MA, USA).

Images were acquired an LSM 700 confocal laser-scanning microscope equipped with a Plan-Neofluar 40× objective lens (numerical aperture = 0.75; both from Carl Zeiss, Oberkochen, Germany) with a pinhole setting that corresponded to a focal plane thickness of less than 1 μm to obtain images of the stained sections. Quantitative analysis was performed as reported previously [16]. To exclude false-positives due to overlapping signal from different cells, randomly selected positive cells were analyzed by scanning the entire z-axis of each cell. Cells were counted under the live mode of confocal scanning.

Calbindin (CB) fluorescence intensity of the DG was measured by using ImageJ.

For positions of doublecortin-positive (DCX+) cells within the GCL, the positive cell positions were expressed by a relative value between the bottom of GCL (0) and the border with the molecular layer (1).

Data were analyzed using the t-test or 2-way ANOVA. Error bars represent standard error of the mean (SEM).

### Antibodies and reagents

We used the following primary antibodies: mouse monoclonal anti-PV (1:2000, Sigma-Aldrich), CB (1:2000, Sigma-Aldrich), calretinin (CR; 1:10000, Millipore, Billerica, MA, USA), and glutamate decarboxylase 67 (GAD67) (1:10,000, Millipore); rat monoclonal anti-BrdU (1:100; Abcam, Cambridge, MA, USA); rabbit polyclonal anti-gamma-aminobutyric acid (GABA; 1:1000, Sigma-Aldrich) and anti-Ki67 (1□:□10, Ylem, Rome, Italy), and neuropeptide Y (NPY, 1□:□2000, Sigma-Aldrich); and goat polyclonal anti-DCX (1:200, Santa Cruz Biotechnology, Dallas, TX, USA). We also used the following secondary antibodies: Alexa Fluor 488 and 594 goat anti-mouse IgG (both 1:200, Life Technologies, Carlsbad, CA, USA), Cy3 goat anti-mouse IgM (1:200, Millipore), and Alexa Fluor 594 goat antirabbit IgG and anti-rat IgG (both 1:200, Life Technologies). Biotinylated *Wisteria floribunda* agglutinin (1:200, Sigma-Aldrich), followed by Alexa Fluor 488 conjugated to streptavidin (10 μg/ml), was used to label PNNs using the method described above [12].

## Results

### Granule cell dematuration without increased adult neurogenesis in the FLX-treated dentate gyrus

We examined the effects of FLX treatment on neuronal maturity and adult neurogenesis in the hippocampus. To evaluate the maturity of hippocampal granule cells, we used DCX to label late-stage progenitor cells and immature granule cells, CR to label immature granule cells, and CB to label mature granule cells. Unexpectedly, CB fluorescence intensity in the DG was unaffected by FLX treatment (P = 0.81; Figure. 1c, d). However, the number of CR-positive (+) cells was significantly increased by FLX treatment (P = 0.043; Figure. 1a, b). A subset of CR+ cells was located within the granule cell layer (GCL) in FLX-treated animals (Figure 1a). We also observed a significant increase in DCX+ immature granule cells in FLX-treated animals (P < 0.001; Figure 2), many of which were located within the GCL, similar to the pattern of CR+ cells (Figure 2b).

**Figure 1.**
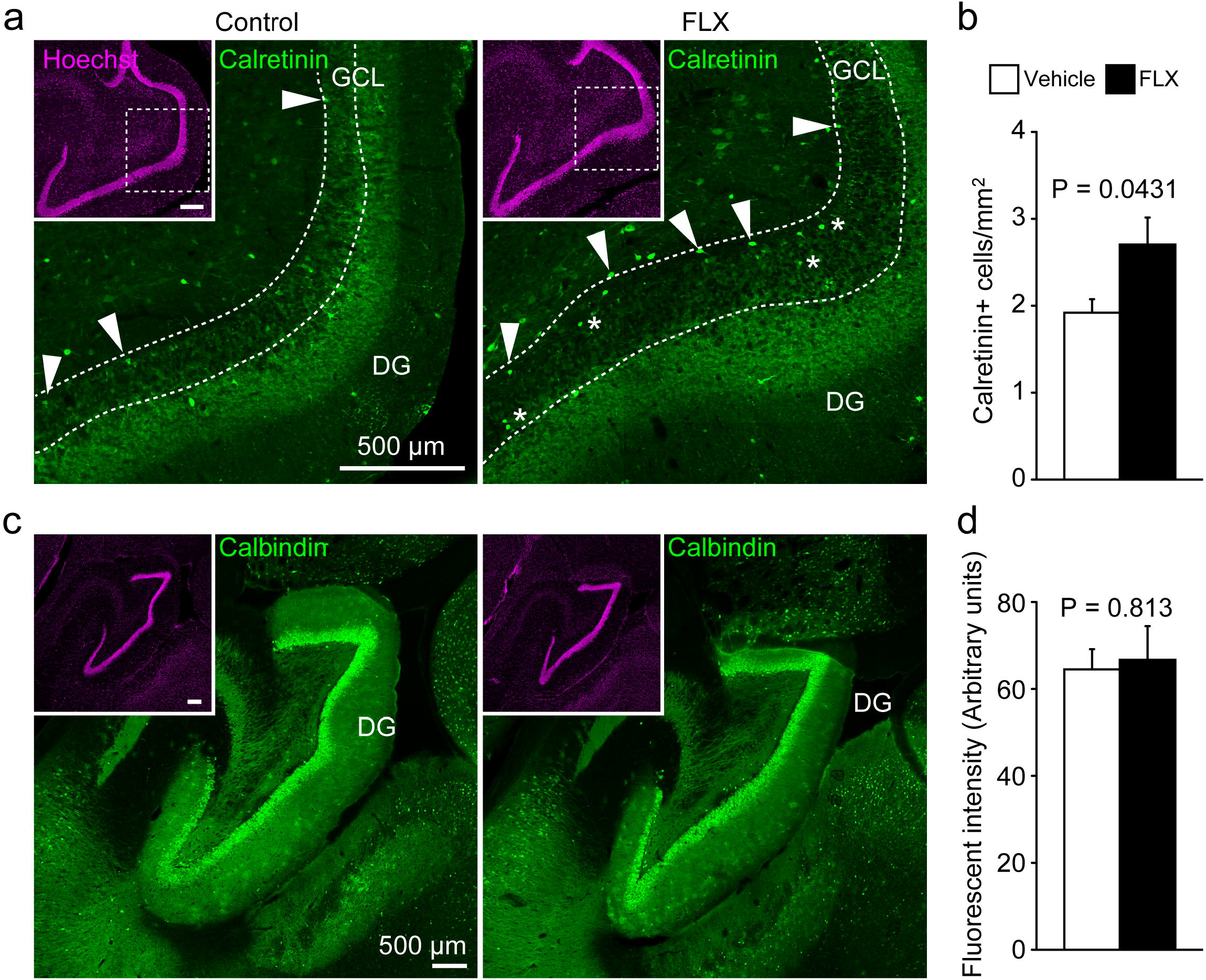
Increased numbers of calretinin-positive cells in the dentate gyrus of fluoxetine-treated marmosets. (A) Representative images of calretinin-positive (CR+) cells in the dentate gyrus (DG) of control (left) and fluoxetine (FLX)-treated marmosets (right). Arrowheads indicate CR+ cells at the base of the granule cell layer (GCL). Asterisks indicate CR+ cells located within the GCL. (B) Quantification of the numbers of CR+ cells. (C) Images of calbindin-positive (CB+) cells in the DG of control (left) and FLX-treated marmosets (right). (D) Quantification of CB fluorescence intensity in the DG.

**Figure 2.**
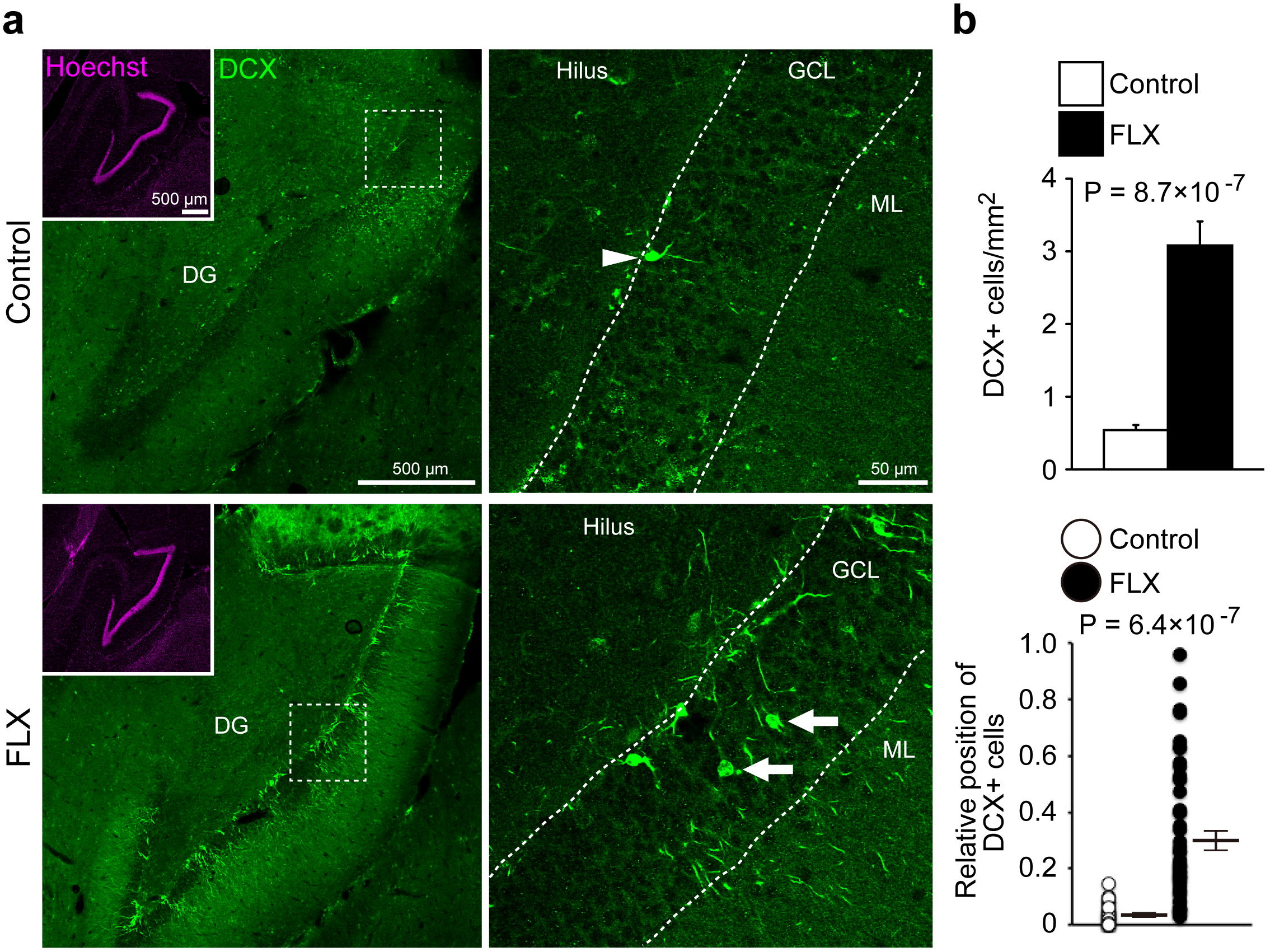
Increased numbers of doublecortin-positive cells in the dentate gyrus of fluoxetine-treated marmosets. (A) Representative images of doublecortin-positive (DCX+) cells in the dentate gyrus (DG) of control (upper row) and fluoxetine (FLX)-treated marmosets (lower row). In the DG of control marmosets, DCX+ cells were located at the base of the granule cell layer (GCL; arrowhead), whereas DCX+ cells were also found throughout the GCL in FLX-treated animals (arrows). (B) Quantification of the numbers and positions of DCX+ cells in the DG. ML, molecular layer.

The increased number of immature granule cells in the FLX-treated GCL raised the possibility of an increase in adult neurogenesis in the DG. To test this, we assessed the effect of FLX on hippocampal adult neurogenesis using BrdU labeling. Although enhancement of adult hippocampal neurogenesis by FLX treatment is a well-known phenomenon in the rodent brain, we did not observe significant increases in adult hippocampal neurogenesis in FLX-treated marmosets compared to controls (P = 0.30; Additional file 1: Figure S1). In both control and FLX-treated marmosets, BrdU+ granule cells were located in the subgranular zone, but not within the GCL (P = 0.60).

### Specific reduction of PV expression and PNNs in the hippocampus

Chronic FLX treatment has been reported to reverse the maturation status of fast-spiking interneurons in the adult rodent hippocampus [11,12]. Thus, we next examined whether FLX treatment alters PV expression or the presence of PNNs, markers of mature fast-spiking interneurons, in the adult marmoset hippocampus. In the DG, the numbers of PV+, PNN+, and PV+/PNN+ cells were significantly decreased by FLX treatment (all P < 0.01; Figure 3a, b). In the CA3 region, the numbers of PV+ (P = 0.0019) and PV+/PNN+ cells (P = 0.032), but not the number of PNN+ cells (P = 0.35), were also decreased by FLX treatment. To determine whether these decreases in PV+ and PNN+ cells reflect apoptosis of PV+ cells, we used TUNEL analysis to measure the numbers of apoptotic cells in FLX-treated and control animals. However, our results show that FLX treatment did not induce apoptosis in the hippocampus (Additional file 1: Figure S2).

**Figure 3.**
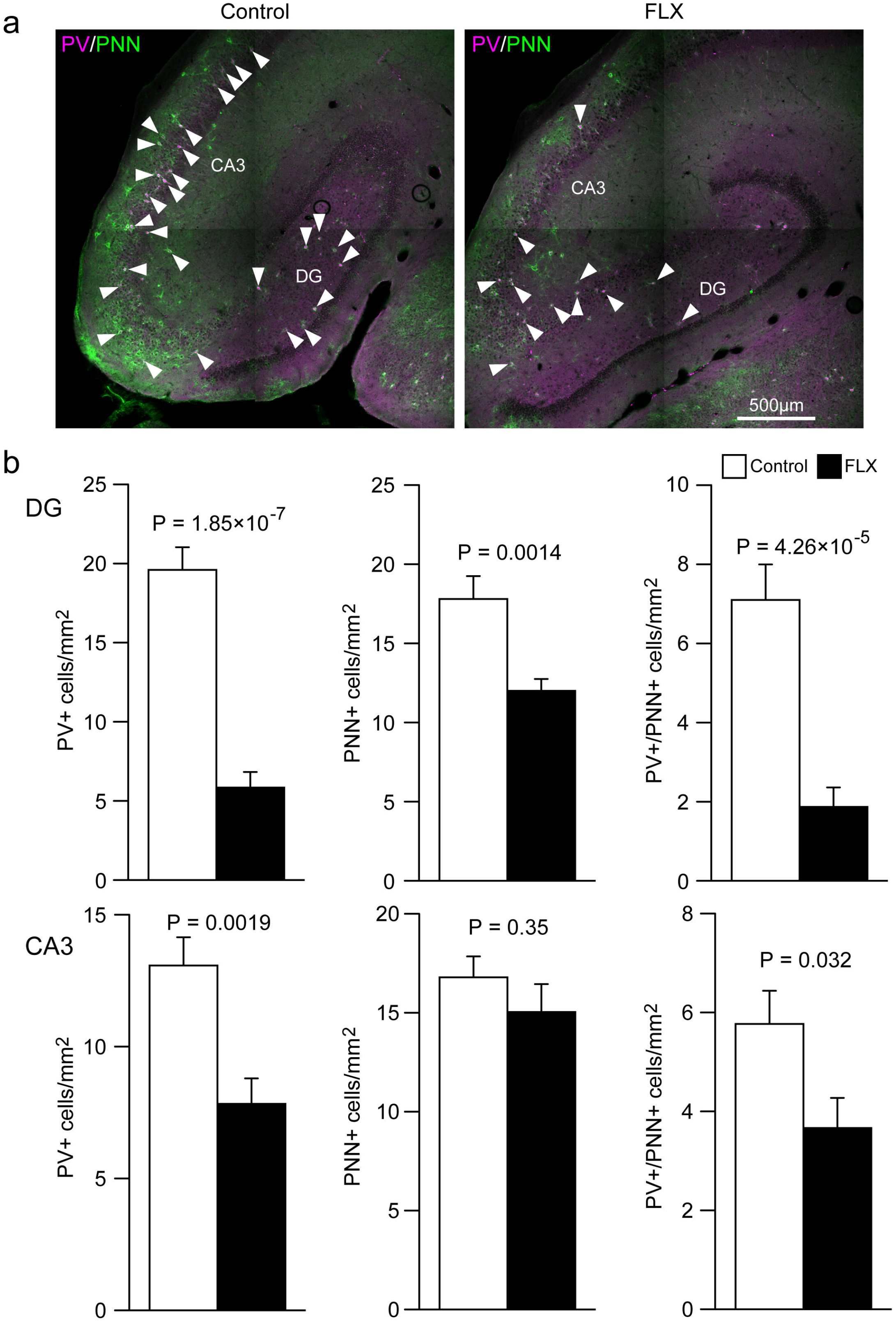
Decreased numbers of parvalbumin and/or perineuronal net-positive cells in the hippocampus of fluoxetine-treated marmosets. (A) Representative images of parvalbumin-positive (PV+; magenta)/perineuronal net-positive (PNN+; □green) cells in the hippocampus of control (left) and fluoxetine (FLX)-treated marmosets (right). (B) Quantification of the numbers of PV+, PNN+, and PV+/PNN+□cells in the dentate gyrus (DG; upper row) and CA3 region (lower row).

Previous studies show that FLX treatment also reverses maturation of fast-spiking interneurons in the amygdala and cerebral cortex of adult mice [11,12]. To determine whether a similar phenomenon occurs in primates, we performed immunostaining for PV and PNNs in the amygdala and cerebral cortex of FLX-treated marmosets. There were no differences between FLX- and control-treated groups in the numbers of PV+, PNN+, or PV+/PNN+ cells in the amygdala or cerebral cortex (all P > 0.05; Additional file 1: Figures S3, S4).

### Increase in adult neurogenesis in the cerebral cortex

In our previous study, we found inhibitory neuron progenitor cells in the cerebral cortex of adult rodents [8], which we termed layer 1 inhibitory neuron progenitor cells (L1-INP cells). Proliferation of L1-INP cells is upregulated by FLX treatment in mice [9]. We first confirmed the existence of L1-INP cells in the cerebral cortex of adult marmosets using the cell markers, GAD67 and Ki67, which can co-label L1-INP cells specifically (Figure 4a). Next, we examined the effects of FLX on cortical neurogenesis in adult marmosets. We observed a non-significant increase in the number of L1-INP cells in the cortex of FLX-treated marmosets compared to controls (P = 0.098; Figure 4b). The number of BrdU+/GAD67+ cells, representing new interneurons, was significantly increased by FLX treatment in the cerebral cortex (P < 0.001; Figure 5). Finally, we investigated the subtypes of newly generated interneurons using the interneuron markers PV, CB, CR, and NPY. A subset of newly generated cells expressed CR (Table 1, Additional file 1: Figure 5); however none of these newly generated cells expressed PV, CB, or NPY (Table 1; Additional file: Figure 6), suggesting that FLX treatment may stimulate the generation or development of specific interneuron subtypes.

**Figure 4.**
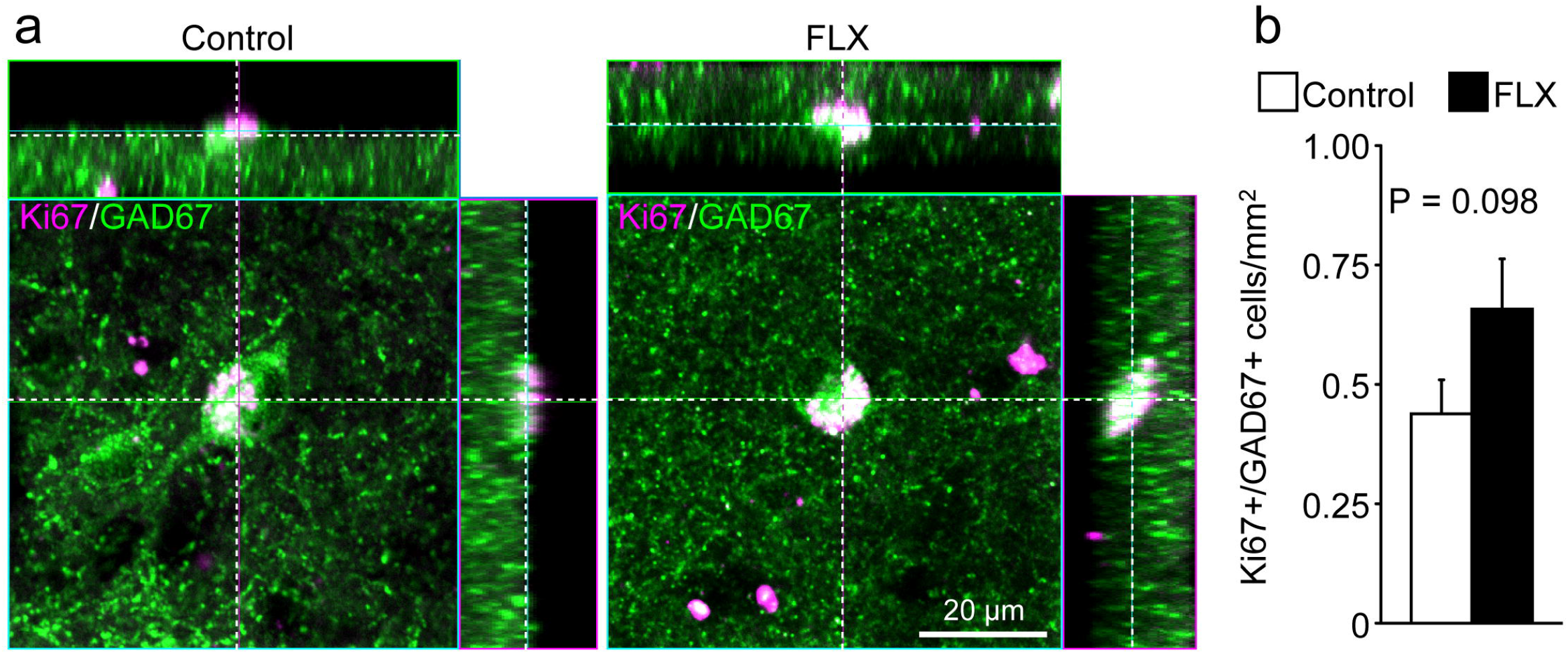
Trending increase in the number of layer 1 inhibitory neuron progenitor cells in the cerebral cortex of fluoxetine-treated marmosets. (A) Representative images of layer 1 inhibitory neuron progenitor (L1-INP) cells in control (left) and fluoxetine (FLX)-treated marmosets (right). Ki67+ (magenta)/glutamate decarboxylase (GAD)67+ (green) cells were identified as L1-INP cells. (B) Quantification of the numbers of L1-INP cells.

**Figure 5.**
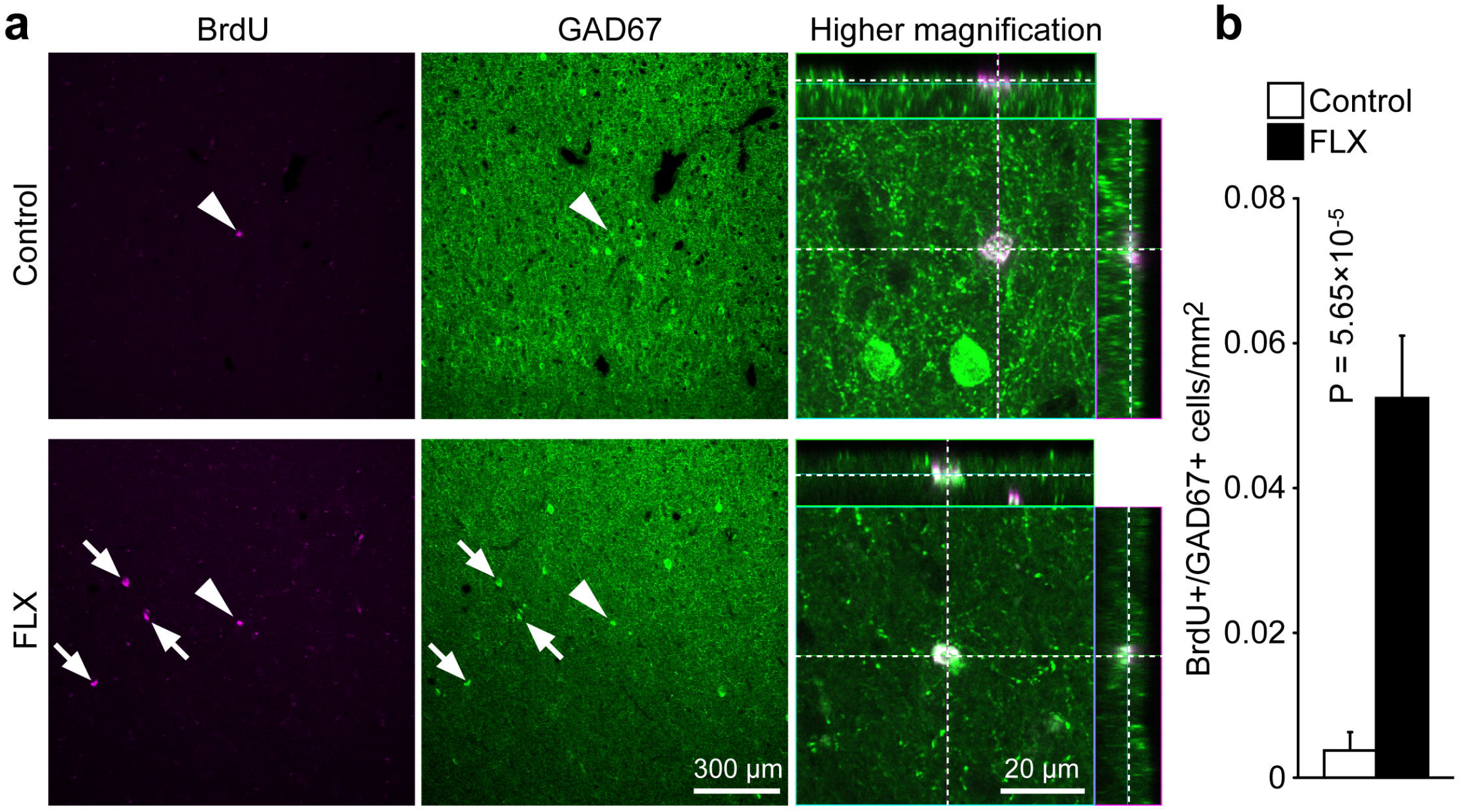
Fluoxetine-induced production of new interneurons in the marmoset cerebral cortex. (A) Newly generated interneurons were labeled with anti-bromodeoxyuridine (BrdU; magenta) and anti-GAD67 antibodies (green). Arrowheads indicate new interneurons whose images are shown at higher magnification. Arrows indicate BrdU+/GAD67+ cells. (B) Quantification of the numbers of newly generated interneurons in the medial frontal cortex of marmosets. Fluoxetine (FLX) treatment significantly increased the generation of new interneurons.

**Table 1.** Subtypes of newly generated interneurons

|  |  | Control | FLX-treated |
| --- | --- | --- | --- |
| CR | BrdU+ | $0.366 \pm 0.208 \text{ cells/mm}^2$ | $3.82 \pm 0.516^{*1}$ |
| | BrdU+/CR+ | $0.122 \pm 0.0814$ | $1.32 \pm 0.223^{*1}$ |
| CB | BrdU+ | $0.475 \pm 0.170$ | $2.44 \pm 0.322^{*1}$ |
|  | BrdU+/CB+ | 0 | 0 |
| PV | BrdU+ | $1.36 \pm 0.325$ | $3.85 \pm 0.596^{*2}$ |
|  | BrdU+/PV+ | 0 | 0 |
| NPY | BrdU+ | $0.610 \pm 0.228$ | $5.56 \pm 0.671^{*1}$ |
|  | BrdU+/NPY+ | 0 | 0 |

Data are presented as mean ± standard error of the mean (SEM) of 3 animals per group. FLX, fluoxetine; CR, calretinin; CB, calbindin; PV, parvalbumin; NPY, neuropeptide Y; BrdU, bromodeoxyuridine. $*^1P < 0.0001$ ; $*^2P = 0.0012$

## Discussion

In this study, we examined the effects of FLX on the brains of adult marmosets. We found that FLX treatment increased the expression of the immature granule cell markers CR and DCX in the DG without enhancement of adult hippocampal neurogenesis. In contrast, PV expression and PNNs, markers of mature fast-spiking interneurons, were significantly decreased in the hippocampus, but not in the cerebral cortex or amygdala, of FLX-treated marmosets compared to controls. In addition, FLX markedly increased the number of BrdU+/GAD67+ cells in the FLX-treated cortex.

CR+ and DCX+ cells were located throughout the GCL in FLX-treated animals but were observed in the deepest portion of the GCL of control animals. Moreover, adult hippocampal neurogenesis, as assessed by BrdU labeling, was not upregulated by FLX in marmosets in this study. Newly generated BrdU+ cells were found in the subgranular zone in both control and FLX-treated marmosets. Taken together, these data suggest that FLX treatment converts exciting mature granule cells to a pseudo-immature state in adult marmosets, which is consistent with previous findings in rodent models (Table 2) [10,15,17]. It remains unclear whether dematuration of dentate granule cells and certain types of hippocampal interneurons is related to the antidepressant and/or adverse behavioral effects of FLX. Interestingly, pseudo-immature molecular and/or electrophysiological features have been observed in certain rodent models of neuropsychiatric disorders such as schizophrenia, bipolar disorder, epilepsy and dementia [18–24]. Dematuration of dentate granule cells is also found in patients with schizophrenia and bipolar disorder [25]. These findings suggest that dematuration of granule cells and hippocampal inhibitory interneurons may be associated with the pathophysiology of some neuropsychiatric disorders. Recent findings suggest that impairments in information processing within neural networks, rather than a chemical imbalance, may be a key mechanism underlying depression [26,27]. Antidepressant-induced dematuration of dentate granule cells and certain types of interneurons in the hippocampus would alter neuronal excitability, morphology, and connectivity, which could gradually improve neuronal information processing and contribute to mood recovery. Consistent with this theory, FLX treatment has been reported to reinstate neural plasticity [28] and promote electrophysiological and functional recovery in the visual cortex of adult amblyopic rats [29]. Furthermore FLX treatment increases expression of brain-derived neurotrophic factor in the brain [30,31]; overexpression of brain-derived neurotrophic factor accelerates PV+ cell maturation, which reduces the capacity for cortical neural plasticity [32,33]. Therefore, dematuration of hippocampal granule cells and PV+ interneurons could reverse losses in synaptic plasticity that occur with age and development, potentially contributing to the antidepressant effects of FLX. Further studies will be required to investigate a possible causal relationship between granule cell and PV+ neuron dematuration and enhanced neural plasticity.

**Table 2.**
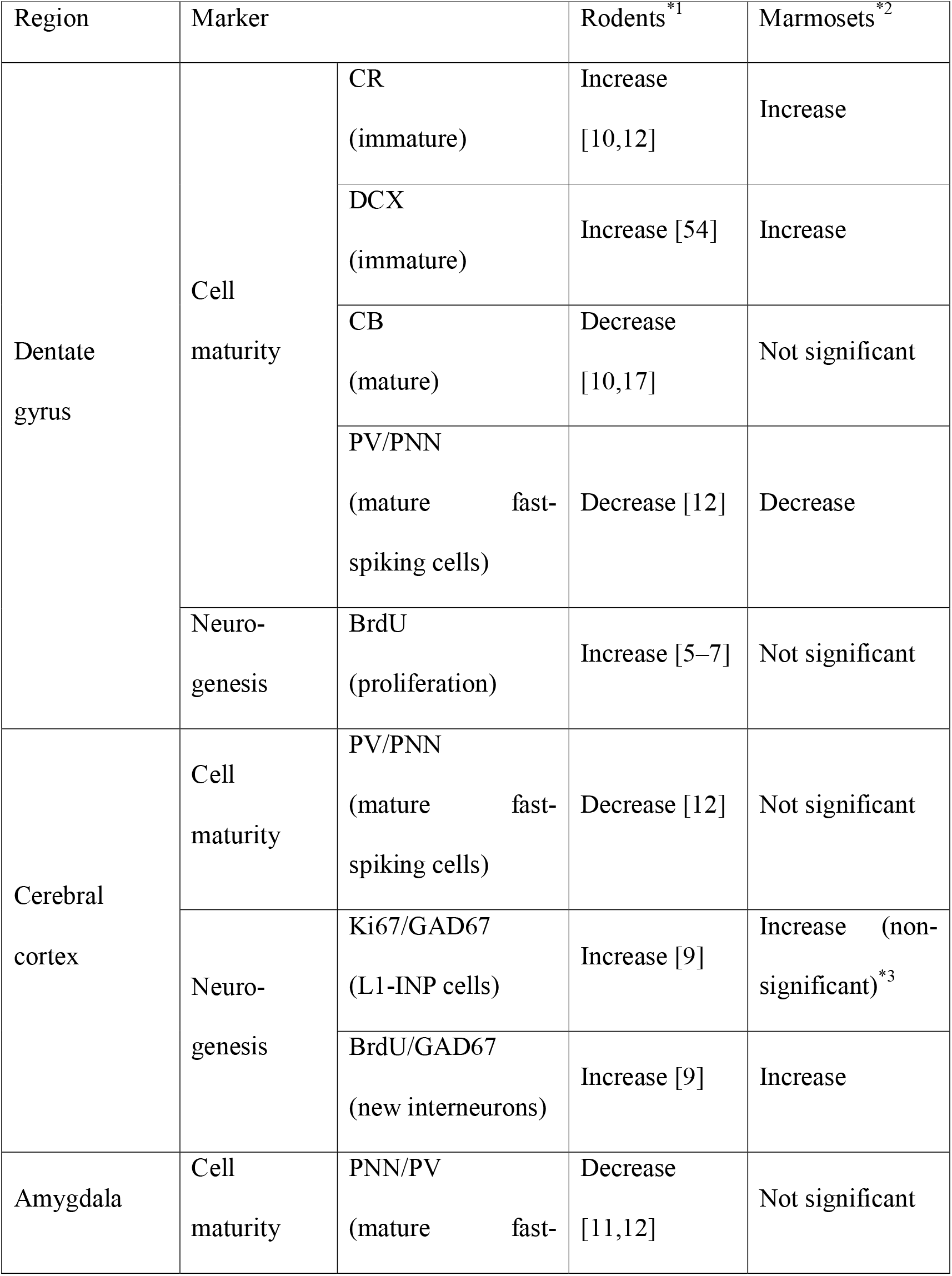
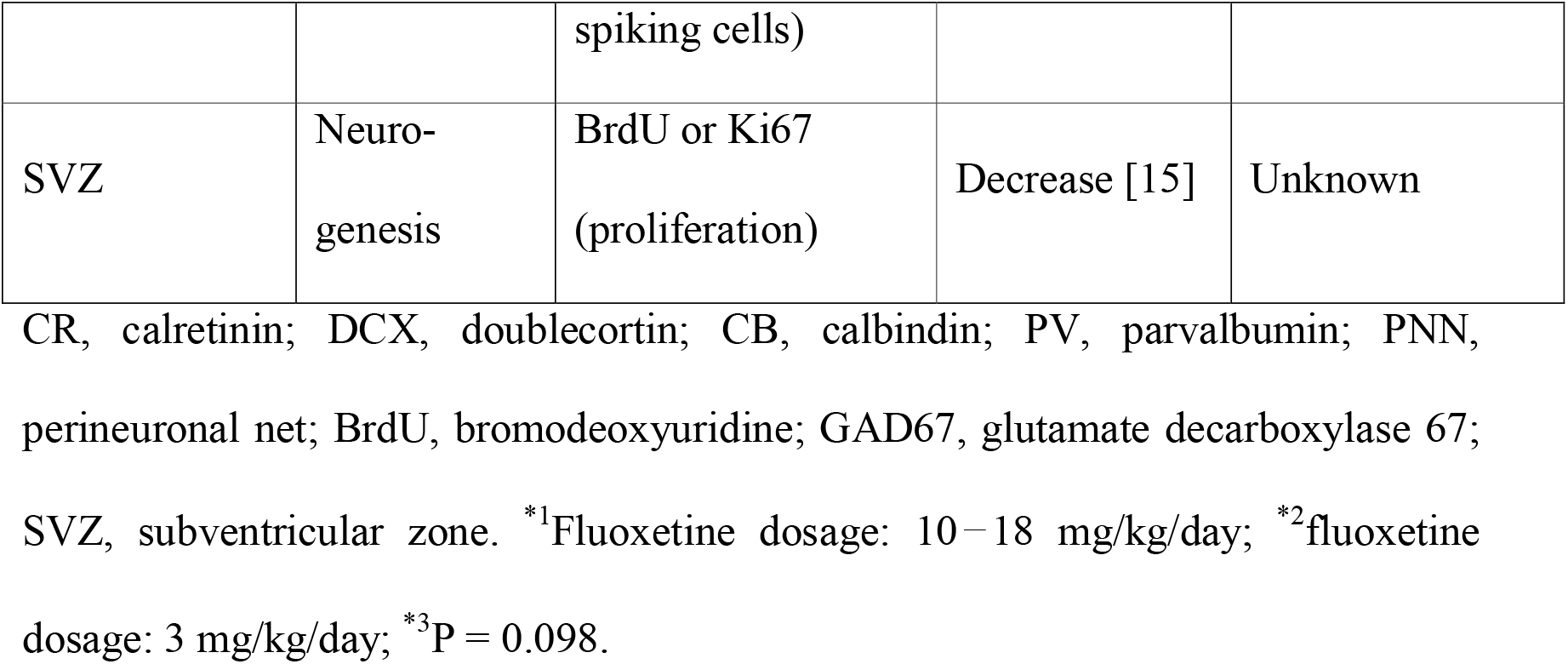
Comparison of fluoxetine-induced effects on cell maturity and neurogenesis in rodents and marmosets

| Region | Marker |  | Rodents<sup>*1</sup> | Marmosets<sup>*2</sup> |
| --- | --- | --- | --- | --- |
| Dentate gyrus | Cell maturity | CR<br>(immature) | Increase<br>[10,12] | Increase |
|  |  | DCX<br>(immature) | Increase [54] | Increase |
|  |  | CB<br>(mature) | Decrease<br>[10,17] | Not significant |
|  |  | PV/PNN<br>(mature fast-spiking cells) | Decrease [12] | Decrease |
|  | Neuro-genesis | BrdU<br>(proliferation) | Increase [5–7] | Not significant |
| Cerebral cortex | Cell maturity | PV/PNN<br>(mature fast-spiking cells) | Decrease [12] | Not significant |
|  | Neuro-genesis | Ki67/GAD67<br>(L1-INP cells) | Increase [9] | Increase (non-significant)<sup>*3</sup> |
|  |  | BrdU/GAD67<br>(new interneurons) | Increase [9] | Increase |
| Amygdala | Cell maturity | PNN/PV<br>(mature fast- spiking cells) | Decrease<br>[11,12] | Not significant |
| SVZ | Neuro-<br>genesis | BrdU or Ki67<br>(proliferation) | Decrease [15] | Unknown |

CR, calretinin; DCX, doublecortin; CB, calbindin; PV, parvalbumin; PNN, perineuronal net; BrdU, bromodeoxyuridine; GAD67, glutamate decarboxylase 67; SVZ, subventricular zone. <sup>\*1</sup>Fluoxetine dosage: 10–18 mg/kg/day; <sup>\*2</sup>fluoxetine dosage: 3 mg/kg/day; <sup>\*3</sup>P = 0.098.

In this study, we did not observe FLX-induced dematuration of PV+ fast-spiking interneurons in the marmoset cortex. A previous study has reported that, while extracellular serotonin concentration in the hippocampus of macaque monkeys remains consistent throughout 3 weeks of FLX treatment, cortical serotonin concentration is increased after 1 week of FLX administration but diminishes thereafter and is no longer significantly elevated at the end of the 3-week treatment period [34]. Thus, effects of FLX in the cortex may dissipate during chronic FLX treatment. If a similar phenomenon occurs in marmosets, this may explain why dematuration of certain cells was not observed in the cortex.

Increased adult hippocampal neurogenesis has been observed in rodents treated with an FLX dosage greater than 10 mg/kg/day (Table 2). In this study, we found that FLX treatment at 3 mg/kg/day did not increase hippocampal neurogenesis in adult marmosets. This dosage was chosen based on a previous report of behavioral alterations in marmosets treated using the same dose [14]; however, it is not clear whether these behavioral changes were due to adult neurogenesis. In other primates such as macaque monkeys and baboons, FLX treatments between 1 and 5 mg/kg/day have been reported to increase adult hippocampal neurogenesis [35,36]. FLX dosages in patients with depression range from 20 to 80 mg daily, which is equivalent to 0.33 to 1.33 mg/kg/day in a human weighing 60 kg; these doses have been shown to upregulate adult hippocampal neurogenesis in humans [37,38]. Marmosets are classified as Platyrrhini (New World monkeys), whereas macaque monkeys and humans are classified as Catarrhini (Old World monkeys and apes). It is possible that more than 3 mg/kg/day of FLX is needed to increase adult hippocampal neurogenesis in marmosets. Thus, it will be important to determine the dose dependency of adult hippocampal neurogenesis in marmosets in future studies.

The present study represents the first demonstration of L1-INP neuronal progenitor cells, identified by Ki67 and GAD67 expression, in cortical layer 1 of the primate brain. In the rodent cortex, FLX increases adult production of GABAergic interneurons from L1-INP cells [9]. We found that FLX treatment also stimulates the production of new GABAergic interneurons in the marmoset cortex (Table 2). These new interneurons express CR, but not CB, PV, or NPY. FLX treatment [9] and ischemia [8] have been shown to induce the production of CR+ and NPY+ cells from L1-INP cells in the rodent cerebral cortex. Although FLX treatment tended to increase the number of L1-INP cells (P = 0.098), it is possible that these effects of FLX in the marmoset cortex had declined after 1 week of treatment because of the failure of FLX treatment to maintain elevated cortical serotonin concentrations, as described above. Thus, it is conceivable that L1-INP cells produce new interneurons during the first week of FLX treatment. Subsequently, new interneurons might remain in the cortex, while the number of L1-INP cells is reduced to control levels. New neurons could also arise from sources other than L1-INP cells, including gray matter, white matter, and the subventricular zone [39]. Further studies will be needed to identify the sources of newly generated interneurons.

CR+ interneurons participate in the disinhibition of other cortical GABAergic interneurons [40–45]. The targets of CR+ cells are mainly Martinotti cells; in turn, the axonal arbors of Martinotti cells form inhibitory contacts with the distal tuft dendrites of pyramidal neurons. Thus, activation of CR+ interneurons leads to disinhibition of the apical dendrites of pyramidal cells, acting as a “gate-opening” mechanism for neural information [46]. A recent study has reported that chronic stress, which causes depression-like behaviors, induces dendritic hypertrophy of Martinotti cells in the medial prefrontal cortex (mPFC) of adult mice [47]. Because decreased mPFC activity has been reported in patients with depression [48], it is possible that pyramidal neuron inhibition may be enhanced by axonal hypertrophy of Martinotti cells in these patients. Thus, the increased number of CR+ interneurons following FLX treatment may reduce the activity of Martinotti cells, which in turn may normalize activity of pyramidal neurons in patients with depression. Previous studies have reported decreases in the number of inhibitory interneurons, amount of GABA, and expression of the GABA-synthesizing enzyme GAD67 in the cerebral cortex of patients with neuropsychiatric disorders such as schizophrenia, depression, dementia, and multiple sclerosis [49–53]. Based on our results, we propose that new GABAergic interneurons of various subtypes may be produced in response to FLX treatment, which compensate for the reduced inhibitory activity in the cerebral cortex of these patients.

In this study, we demonstrated that hippocampal dematuration and cortical neurogenesis occurred in FLX-treated marmosets, similar to previous results obtained using rodent models. However, FLX treatment had little effect on hippocampal neurogenesis and dematuration of certain interneuron subtypes in the marmoset amygdala and cerebral cortex. Further studies will be necessary to determine whether FLX-induced hippocampal neuron dematuration and cortical neurogenesis are involved in the therapeutic mechanisms and/or adverse effects of FLX.

## List of abbreviations

BrdU: bromodeoxyuridine
CA1: cornus ammonis 1
CA3: cornus ammonis 3
GABA: gamma-aminobutyric acid
CB: calbindin
CNS: central nervous system
CR: calretinin
DCX: doublecortin
DG: dentate gyrus
FLX: fluoxetine
GAD67: glutamate decarboxylase 67
GCL: granule cell layer
L1-INP cells: layer 1 inhibitory neuron progenitor cells
mPFC: medial prefrontal cortex
NPY: neuropeptide Y
PBS: phosphate-buffered saline
PNN: perineuronal net
PV: parvalbumin
SSRI: selective serotonin reuptake inhibitor

## Declarations

### Ethics approval and consent to participate

All animal experiments were planned and executed in strict accordance with the Guidelines for Care and Use of Nonhuman Primates (Ver. 3; Primate Research Institute, Kyoto University, 2010). The protocol was approved by the Animal Welfare and Animal Care Committee at the Primate Research Institute of Kyoto University (Permission Nos. 2011-B-8, 2012-B-63, and 2013-B-63).

### Consent for publication

Not applicable

### Availability of data and material

All data supporting this article are available from the corresponding author upon reasonable request.

## Acknowledgements

We thank Rika Takeuchi (currently Chubu University, Aichi, Japan) for her assistance with immunohistological analysis.

## Authors’ contributions

K.O., H.H., and T.M. wrote the manuscript. K.O., K.N., and T.M. were involved in experimental design. K.O., M.M., and R.T. performed most of the experiments. All authors revised the final version of the manuscript.

## Additional files

Additional file 1:

Figure S1. No significant change in the number of bromodeoxyuridine-positive cells in the dentate gyrus of fluoxetine-treated marmosets.

Figure S2. No increase in apoptosis in the cortex and hippocampus of fluoxetine-treated marmosets.

Figure S3. No significant changes in the numbers of parvalbumin and/or perineuronal net-positive cells in the amygdala of fluoxetine-treated marmosets.

Figure S4. No significant changes in the numbers of parvalbumin and/or perineuronal net-positive cells in the cerebral cortex of fluoxetine-treated marmosets.

Figure S5. Production of new calretinin-positive interneurons in the cerebral cortex of fluoxetine-treated marmosets.

Figure S6. No expression of parvalbumin, calbindin, or neuropeptide Y in bromodeoxyuridine-positive cells in the marmoset cerebral cortex.

