## Supplemental Information for "Fluoxetine-induced dematuration of hippocampal neurons and adult cortical neurogenesis in the common marmoset"

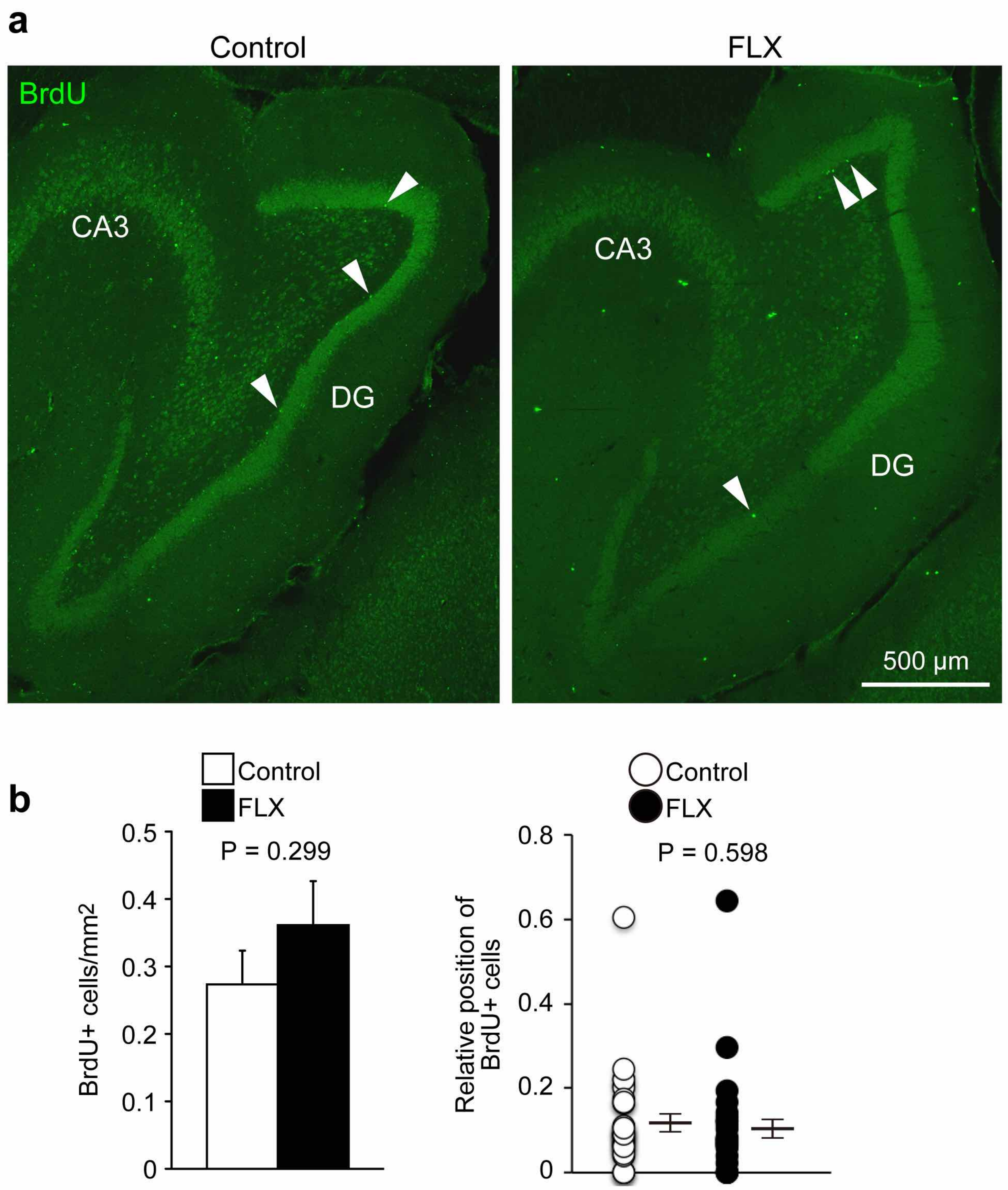


Supplemental Figure S1. No significant change in the number of bromodeoxyuridine-positive cells in the dentate gyrus of fluoxetine-treated marmosets. (a) Representative images of the dentate gyrus (DG) of control and fluoxetine (FLX)-treated marmosets. Arrowheads indicate bromodeoxyuridine-positive (BrdU+) cells. (b) Quantification of the numbers of BrdU+ cells.

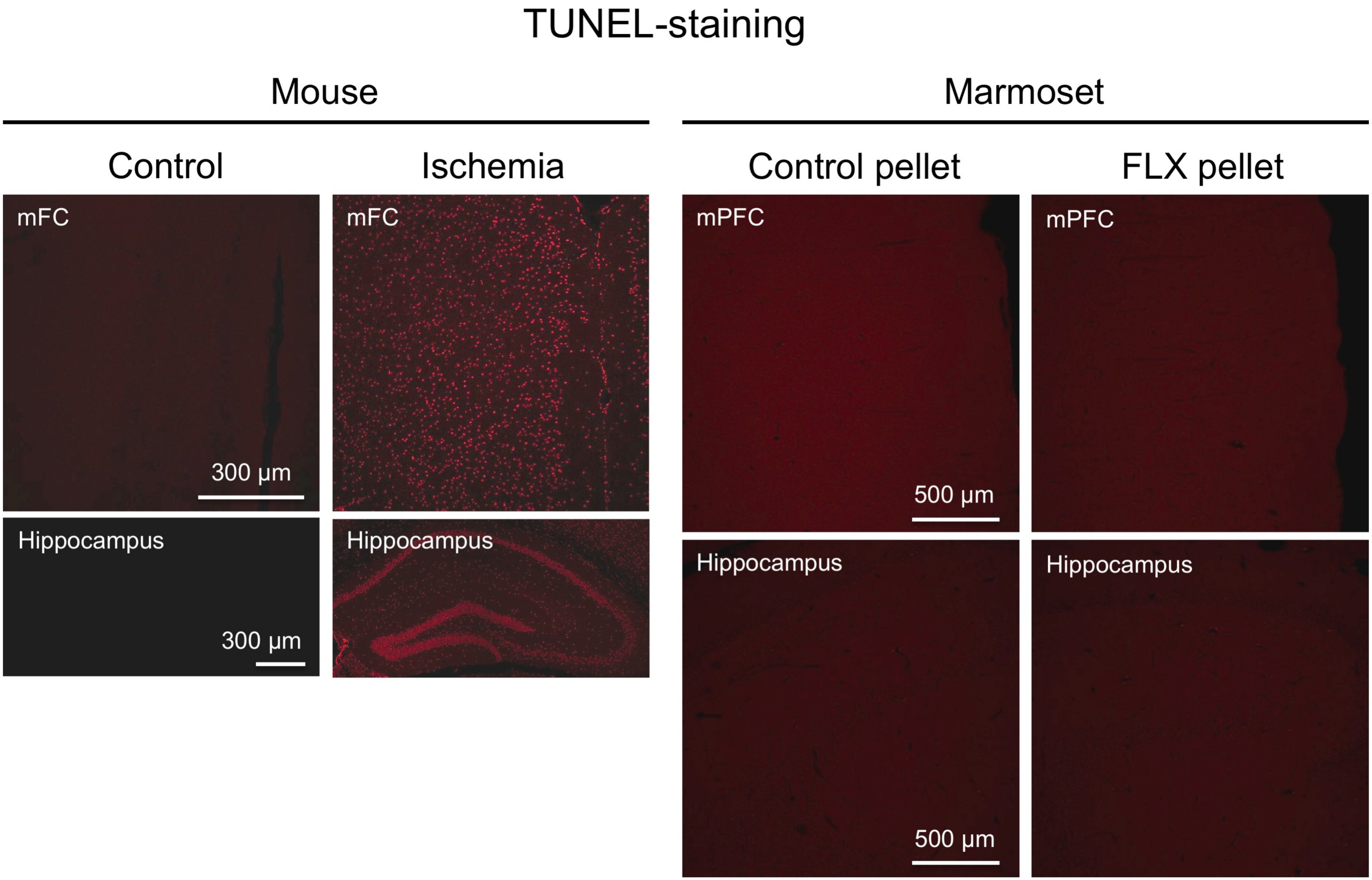


Supplemental Figure S2. No increase in apoptosis in the cortex and hippocampus of fluoxetine-treated marmosets. Ischemic brains of mice were used as a positive control for detection of apoptotic cells (left panels) by terminal deoxynucleotidyl transferase-mediated dUTP nick-end labeling (TUNEL). Few apoptotic cells were observed in the cortex and hippocampus of fluoxetine (FLX)-treated marmosets. mFC, medial frontal cortex; mPFC, medial prefrontal cortex.

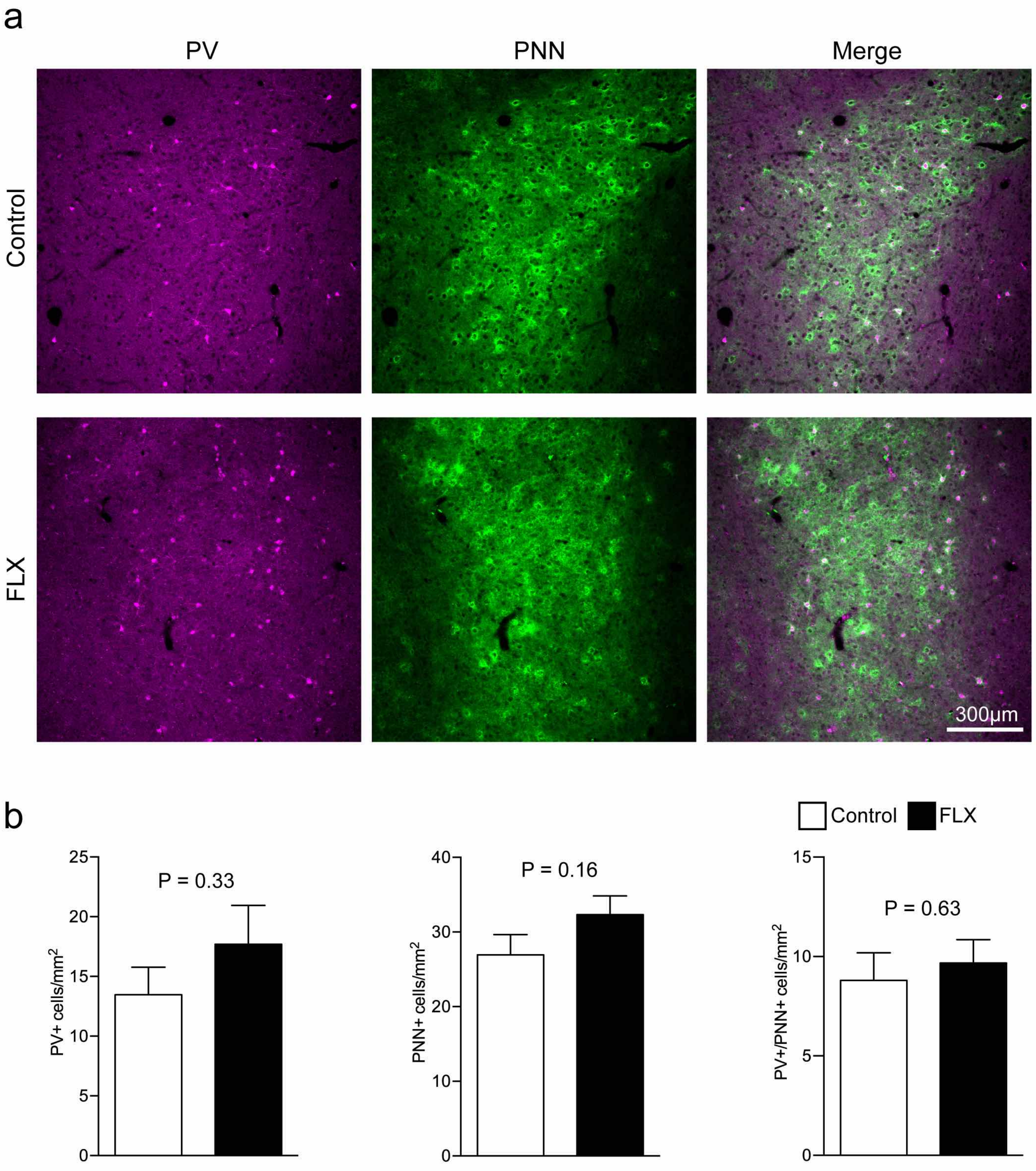


Supplemental Figure S3. No significant changes in the numbers of parvalbumin and/or perineuronal net-positive cells in the amygdala of fluoxetine-treated marmosets. (a) Representative images of parvalbumin-positive (PV+) and perineuronal net-positive (PNN+) cells in the amygdala. (b) Quantification of the numbers of PV+, PNN+, and PV+/PNN+ cells.

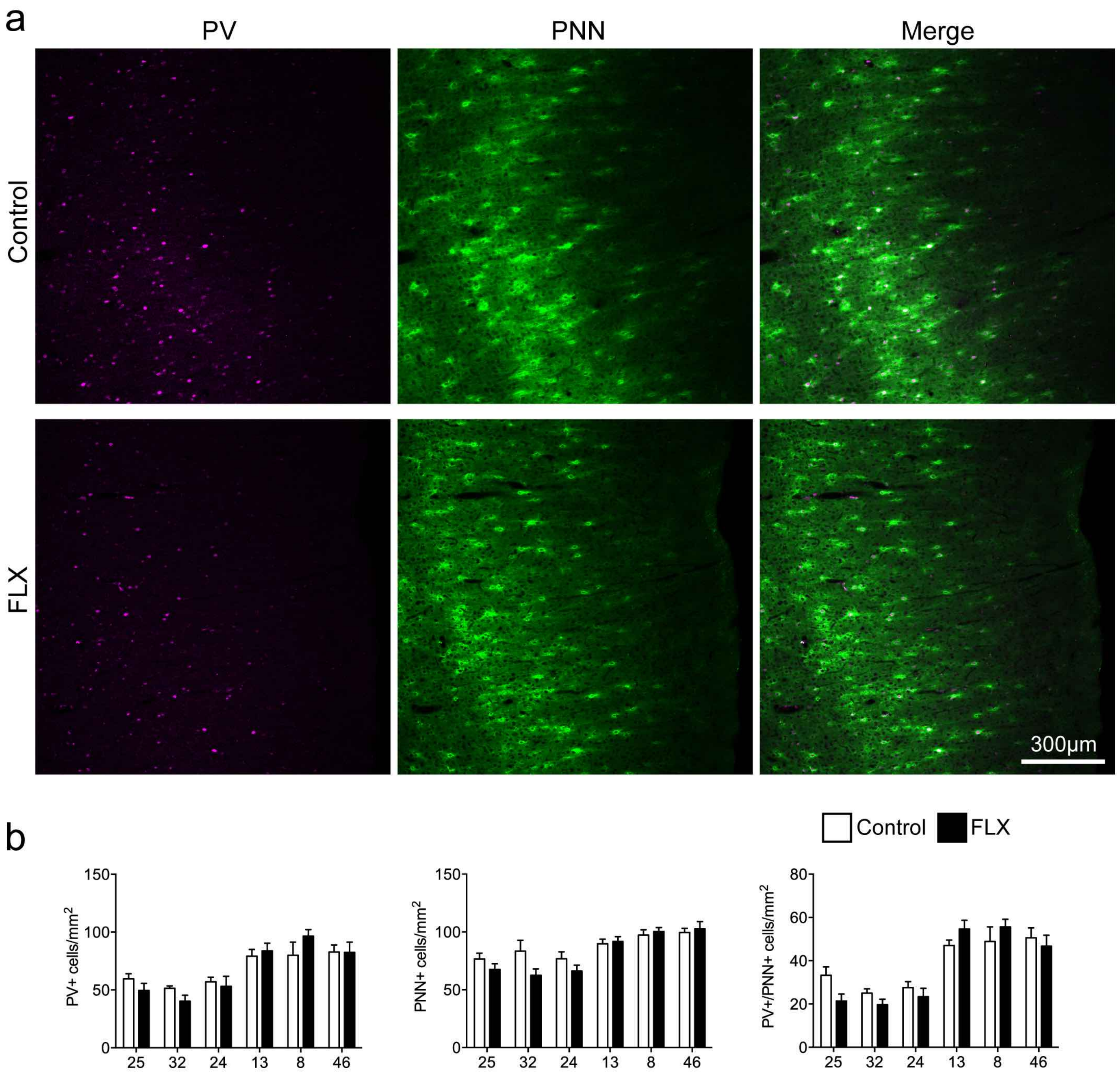


Supplemental Figure S4. No significant changes in the numbers of parvalbumin and/or perineuronal net-positive cells in the cerebral cortex of fluoxetine-treated marmosets. (a) Representative images of parvalbumin-positive (PV+) and perineuronal net-positive (PNN+) cells in area 32 of the cerebral cortex of control (upper row) and fluoxetine (FLX)-treated animals (lower row). (b) Quantification of the numbers of PV+, PNN+, and PV+/PNN+ cells in areas 25, 32, 24, 13, 8, and 46. Data were analyzed by two-way ANOVA.

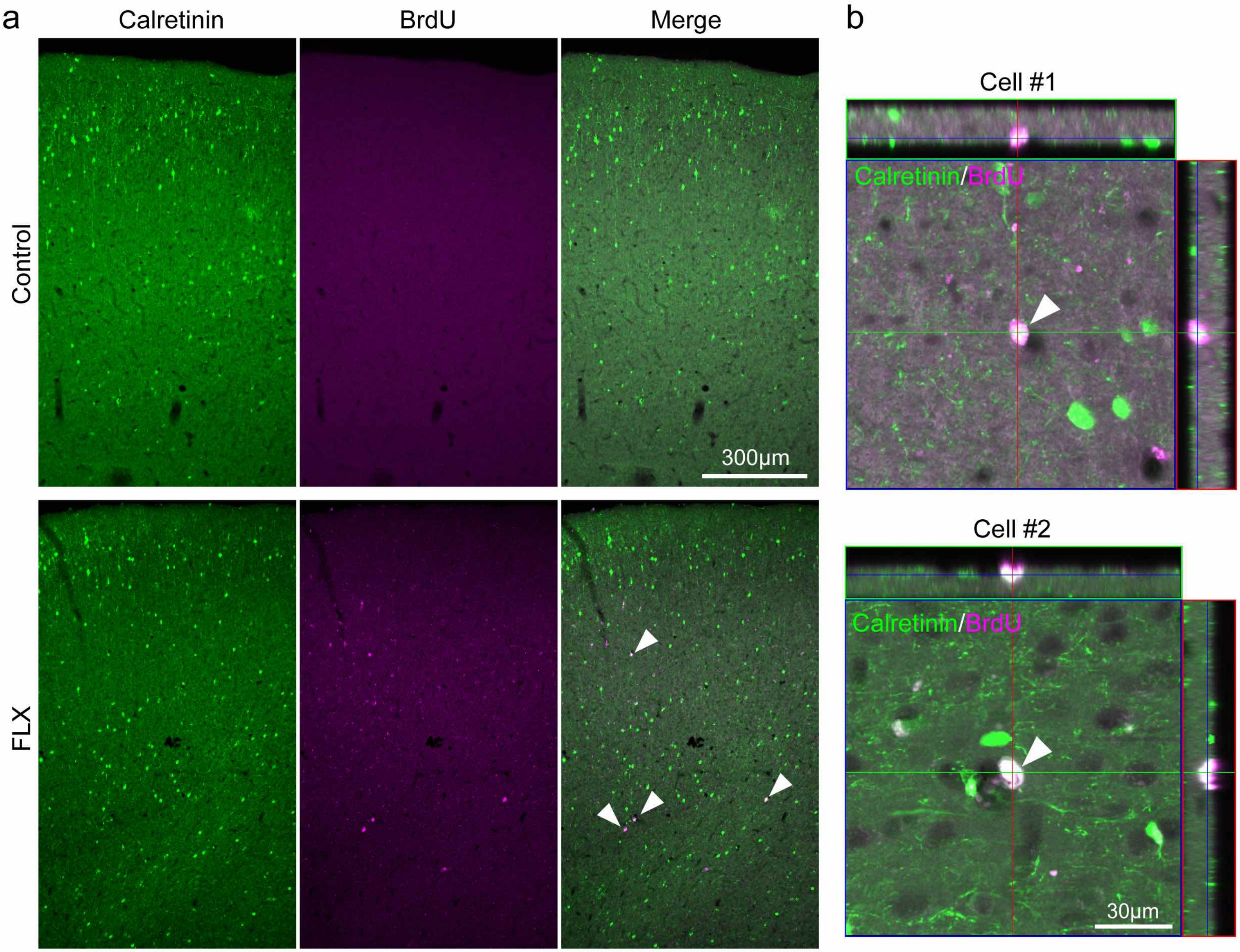


Supplemental Figure S5. Production of new calretinin-positive interneurons in the cerebral cortex of fluoxetine-treated marmosets. (a) Representative images of bromodeoxyuridine-positive (BrdU+)/calretinin-positive (CR+) cells in area 8 of the cerebral cortex of control (upper row) and fluoxetine (FLX)-treated animals (lower row). Arrowheads indicate BrdU+/CR+ cells. (b) Higher-magnification images of BrdU+/CR+ cells in FLX-treated marmosets.

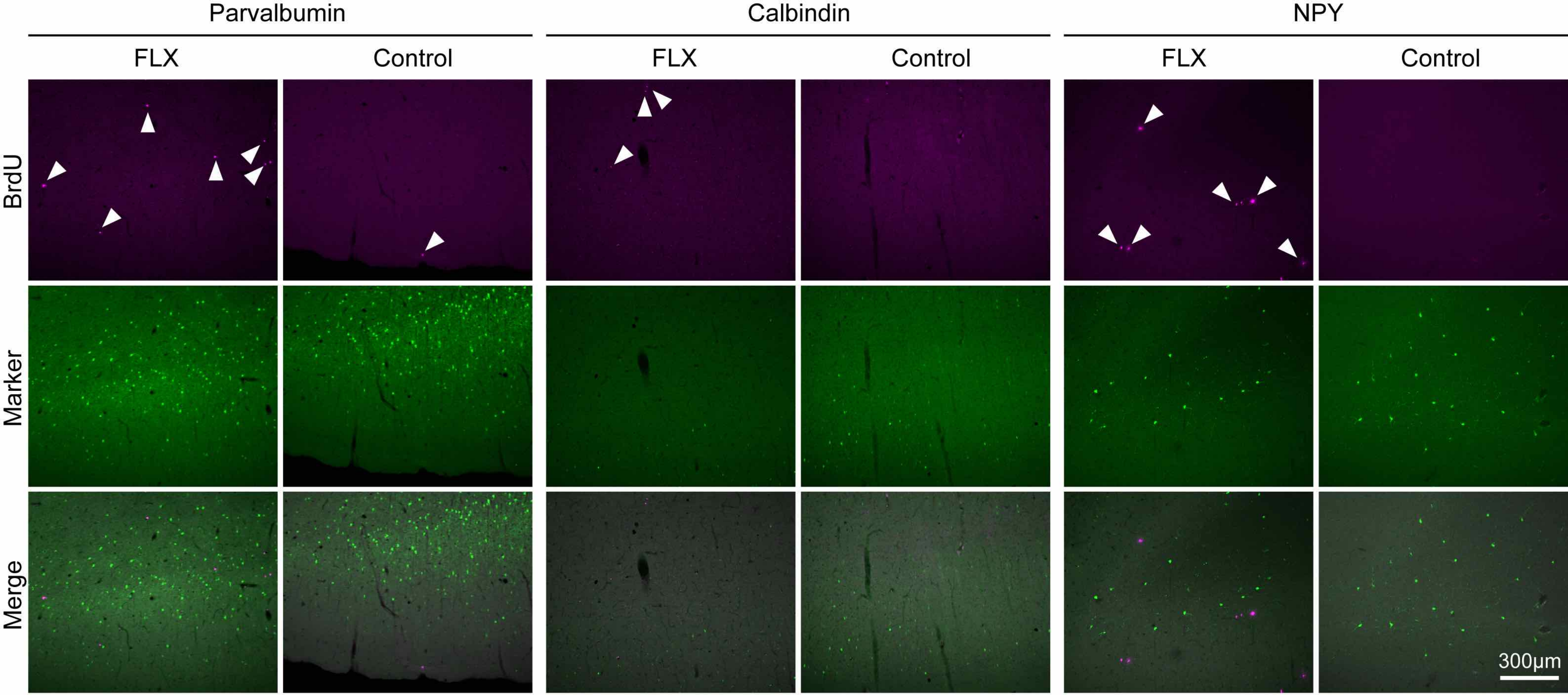


Supplemental Figure S6. No expression of parvalbumin, calbindin, or neuropeptide Y in bromodeoxyuridine-positive cells in the marmoset cerebral cortex. Representative images of bromodeoxyuridine-positive (BrdU+) and neuropeptide Y- (NPY+), calbindin- (CB+), or parvalbumin-positive (PV+) cells in area 8 of the cerebral cortex of control (upper rows) and fluoxetine (FLX)-treated animals (lower rows). Arrowheads indicate bromodeoxyuridine-positive (BrdU+) cells.
